# Rspo2 inhibits TCF3 phosphorylation to antagonize Wnt signaling during vertebrate anteroposterior axis specification

**DOI:** 10.1101/2020.10.30.362541

**Authors:** Alice H. Reis, Sergei Y. Sokol

## Abstract

The Wnt pathway activates target genes by controlling the β-catenin-T-cell factor (TCF) transcriptional complex during embryonic development and cancer. This pathway can be potentiated by R-spondins, a family of proteins that bind RNF43/ZNRF3 E3 ubiquitin ligases and LGR4/5 receptors to prevent Frizzled degradation. Here we demonstrate that, during *Xenopus* anteroposterior axis specification, Rspo2 functions as a Wnt antagonist, both morphologically and at the level of gene targets and pathway mediators. Unexpectedly, the binding to RNF43/ZNRF3 and LGR4/5 was not required for the Wnt inhibitory activity. Moreover, Rspo2 did not influence Dishevelled phosphorylation in response to Wnt ligands, suggesting that Frizzled activity is not affected. Further analysis indicated that the Wnt antagonism is due to the inhibitory effect of Rspo2 on TCF3/TCF7L1 phosphorylation that normally leads to target gene activation. Consistent with this mechanism, Rspo2 anteriorizing activity has been rescued in TCF3-depleted embryos. These observations suggest that Rspo2 is a context-specific regulator of TCF3 phosphorylation and Wnt signaling.

## Introduction

The Wnt pathway is a key conserved developmental pathway that is utilized multiple times during animal development and frequently misregulated in disease (MacDonald et al., 2009; Nusse and Clevers, 2017). Secreted Wnt proteins associate with Frizzled (Fzd) receptors and LRP5/6 coreceptors to stabilize β-catenin and promote β-catenin/T-cell factor (TCF)-dependent transcription. TCF3, also known as TCF7L1, is a predominant embryonic TCF that functions as a transcriptional repressor during early development (Hikasa et al., 2010; Kim et al., 2000; Nguyen et al., 2006). In the presence of Wnt ligands, TCF3 is phosphorylated, followed by its dissociation from target promoters and transcriptional activation that can involve other TCF/LEF transcription factors including TCF1/TCF7 (Cadigan and Waterman, 2012; Hikasa and Sokol, 2011). Whereas many studies of the Wnt pathway mainly focused on the control of β-catenin stability, the regulation of TCF protein activity has been less understood.

R-spondins are prominent extracellular modulators of Wnt signaling in vertebrates (Niehrs, 2012). The R-spondin (Rspo) family consists of four secreted proteins that share high similarity of amino acid sequence and structural organization and play critical roles in development, stem cell biology and cancer (de Lau et al., 2014; Raslan and Yoon, 2019). Mice lacking the *rspo2* gene die at birth due to lung, limb and craniofacial defects, illustrating its essential functions in embryogenesis (Aoki et al., 2008; Bell et al., 2008; Nam et al., 2007; Yamada et al., 2009). Additionally, Rspo2 has been implicated in fish skeletogenesis (Tatsumi et al., 2014) and frog muscle development (Kazanskaya et al., 2004).

The closely related Rspo3 functions in early angiogenesis in mouse and *Xenopus* embryos (Aoki et al., 2007; Kazanskaya et al., 2008). These observations highlight the important functions of R-spondins during embryonic development.

R-spondins are thought to exert their effects by potentiating Wnt/β-catenin signaling (Bell et al., 2008; de Lau et al., 2014; Kazanskaya et al., 2004; Raslan and Yoon, 2019). R-spondins bind LGR4/5 receptors and the E3 ubiquitin ligases ZNRF3/RNF43, thereby stabilizing Frizzled and promoting Wnt signaling (Carmon et al., 2011; de Lau et al., 2011; Hao et al., 2012; Koo et al., 2012). Recent analysis revealed that the mechanisms used by R-spondins to modulate Wnt signaling are more complex (Park et al., 2018; Yan et al., 2017). R-spondins can affect the Wnt pathway independently of LGR4/5 (Lebensohn and Rohatgi, 2018; Szenker-Ravi et al., 2018), indicating the existence of multiple receptors and alternative signaling pathways. Supporting this view, the interaction with heparan sulfate chains is sufficient for R-spondins to modulate Wnt signaling in cells lacking LGR4/5/6 receptors (Dubey et al., 2020).

The Wnt pathway plays crucial and specific roles during anteroposterior axis specification and patterning (De Robertis and Kuroda, 2004; Heasman, 2006; Hikasa and Sokol, 2013). Wnt signals promote posterior structures in the embryo, whereas secreted Wnt antagonists in the anterior region are responsible for head development (Itoh et al., 1995; Kiecker and Niehrs, 2001). One of the reported gain-of-function phenotypes for Rspo2 in *Xenopus* is the formation of ectopic cement gland (Kazanskaya et al., 2004), an anterior mucus-secreting organ (Picard, 1975; Sive et al., 1989). Notably, this phenotype is a common property of Wnt antagonists including GSK3 (Itoh et al., 1995), Axin-related protein (Itoh et al., 2000) and is exhibited in embryos with depleted β-catenin (Heasman et al., 2000). Since this observation is contrary to what is expected of a Wnt coactivator, we decided to reevaluate a role of Rspo2 in the Wnt pathway during *Xenopus* anteroposterior patterning. We show that Rspo2 inhibits Wnt signaling in a manner that is independent of the LGR4/5 and ZNRF3/RNF43 interactions. Mechanistically, we find that Rspo2 downregulates TCF3 phosphorylation that is necessary for target gene activation. Our findings indicate that the same R-spondin can function in a context-dependent manner to either stimulate or inhibit the Wnt pathway.

## Results

### Rspo2 is essential for anterior development

To better characterize the role of Rspo2 in anteroposterior patterning, Rspo2 RNA was injected into early embryos. Confirming earlier findings (Kazanskaya et al., 2004; Reis and Sokol, 2020), the injected embryos developed enlarged cement gland and other head structures (Fig. 1A, B). We next defined early genes induced by Rspo2 by carrying out transcriptome analysis in the ectoderm explants expressing Rspo2 RNA at the onset of gastrulation. We observed the induction of many anterior genes, including *otx1, otx2, and otx5, zic3, rax* (Supplementary Fig. 1). RT-qPCR validated the induction of *otx2* and *ag1* (Sive et al., 1989), whereas the level of *krt12.4*, epidermal keratin, has decreased (Fig. 1C). O*tx* genes are required for anterior development and cement gland formation (Blitz and Cho, 1995; Pannese et al., 1995), suggesting that they could be responsible for the observed Rspo2 activity.

**Figure 1.**
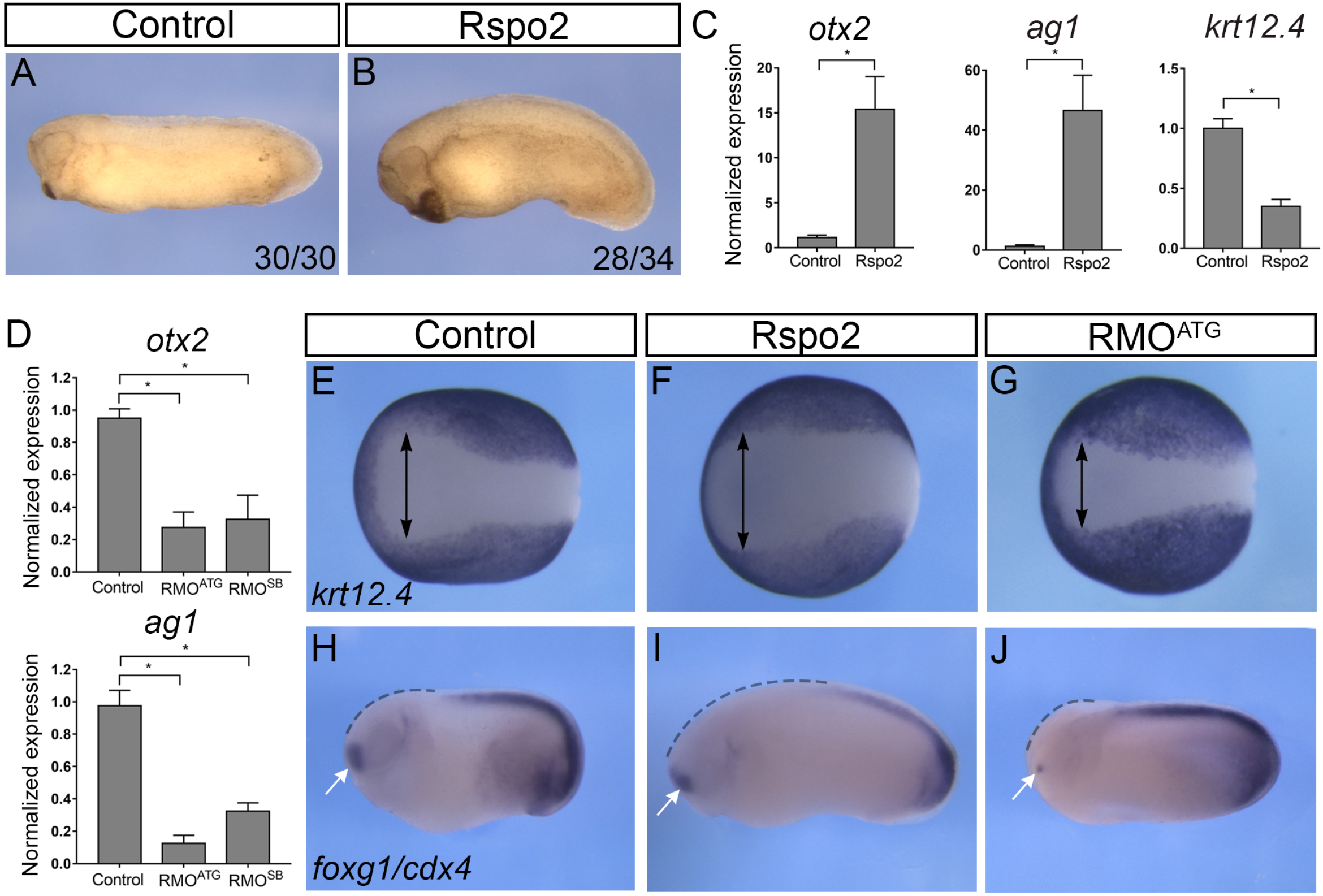
Rspo2 function is essential for anterior development. (A, B) Four-cell embryos were injected with 0.5 ng of Rspo2 RNA into both animal-ventral blastomeres and cultured until stage 28. (A) Uninjected control embryo. (B) Representative embryo injected with Rspo2 RNA. The penetrance is indicated as the ratio of the number of embryos with the phenotype and the total number of injected embryos. (C) The effect of Rspo2 on gene marker expression. Animal pole explants were dissected at stage 9 from embryos overexpressing Rspo2 RNA or uninjected controls. RT-qPCR analysis was carried out for *otx2, ag1*, and *krt12.4* at stage 18. (D) Altered gene expression in Rspo2 morphants. RNA was isolated from stage 18 control embryos or embryos depleted of Rspo2. RT-qPCR for *ag1* and *otx2* was carried out in triplicates. (C, D) Each graph is a single experiment with triplicate samples, representative from at least 3 independent experiments. Means +/- s. d. are shown. Statistical significance has been assessed by Student’s *t* test, *, p<0.05. (E-J) *In situ* hybridization of control and manipulated stage 16 or 25 embryos with *krt12.4* (E-G), *foxg1* and *cdx4* (H-J) probes. (E-G) Width of the anterior neural plate is shown as lack of *krt12.4* (arrows). (H-J) The *foxg1* domain is indicated by white arrows, the anterior region lacking *cdx4* - by dashed lines. See Supplementary Table 1 for quantification.

In complementary experiments, Rspo2 has been depleted using previously characterized translation-blocking (RMO^ATG^) and splicing-blocking (RMO^SB^) morpholino oligonucleotides (MOs)(Reis and Sokol, 2020). Both MOs strongly reduced *otx2* and *ag1* levels (Fig. 1D), causing severe head defects (Reis and Sokol, 2020). Because RMO^ATG^ was more effective for the Rspo2 knockdown, it has been predominantly used in subsequent experiments.

Examination of ectodermal markers in Rspo2-overexpressing embryos by *in situ* hybridization revealed the expansion of the anterior neural plate at the expense of epidermal keratin *krt12.4* in embryos overexpressing Rspo2 (Fig. 1E, F, Supplementary Table 1). By contrast, the anterior neural domain was reduced in Rspo2 morphants (Fig. 1G, Supplementary Table 1). Similarly, the domains of *foxg1* and *cdx4* expression have been coordinately regulated by Rspo2 manipulation (Fig. 1H-J). Taken together, these observations illustrate an essential role of Rspo2 in anterior development.

We next evaluated whether the observed effect of Rspo2 is mediated by its interaction with ZNRF3/RNF43 and LGR4/5 (Carmon et al., 2011; de Lau et al., 2011; Hao et al., 2012; Koo et al., 2012; Wang et al., 2013; Xie et al., 2013; Zebisch and Jones, 2015). We generated point mutations in the sequence of the furin-like domains that eliminate the binding of Rspo2 to ZNRF3/RNF43 and LGR4/5 (Xie et al., 2013). These mutants were expressed at similar levels and induced enlarged or ectopic cement glands in the majority of the injected embryos (Supplementary Fig. 2). These findings suggest that the binding of Znrf3/Rnf43 and Lgr4/5 is not required for Rspo2 ability to anteriorize the embryo.

### Rspo2 is a Wnt antagonist

The anteriorized phenotype caused by Rspo2 is similar to the ones generated by Wnt antagonists (Glinka et al., 1998; Heasman et al., 2000; Itoh et al., 2000; Itoh et al., 1995; Wang et al., 1997; Zhang et al., 2012). We therefore wanted to examine whether Rspo2 could antagonize Wnt signaling.

During gastrulation, Wnt8 enhances posterior development by inducing a distinct set of target genes (Christian and Moon, 1993; Hamilton et al., 2001; Hikasa and Sokol, 2013). To evaluate how Rspo2 affects Wnt signaling, it was co-expressed with Wnt8 in dorsal blastomeres of four-cell embryos. As expected, the majority of embryos injected with *wnt8* DNA became headless (Fig. 2A-C). Separate injections of *Rspo2* RNA into dorsal blastomeres produced blastopore closure defects (Fig. 2D). When coexpressed with Wnt8, Rspo2 completely rescued the headless phenotype in most of the injected embryos (Fig. 2E, F), revealing its Wnt inhibitory activity.

**Figure 2.**
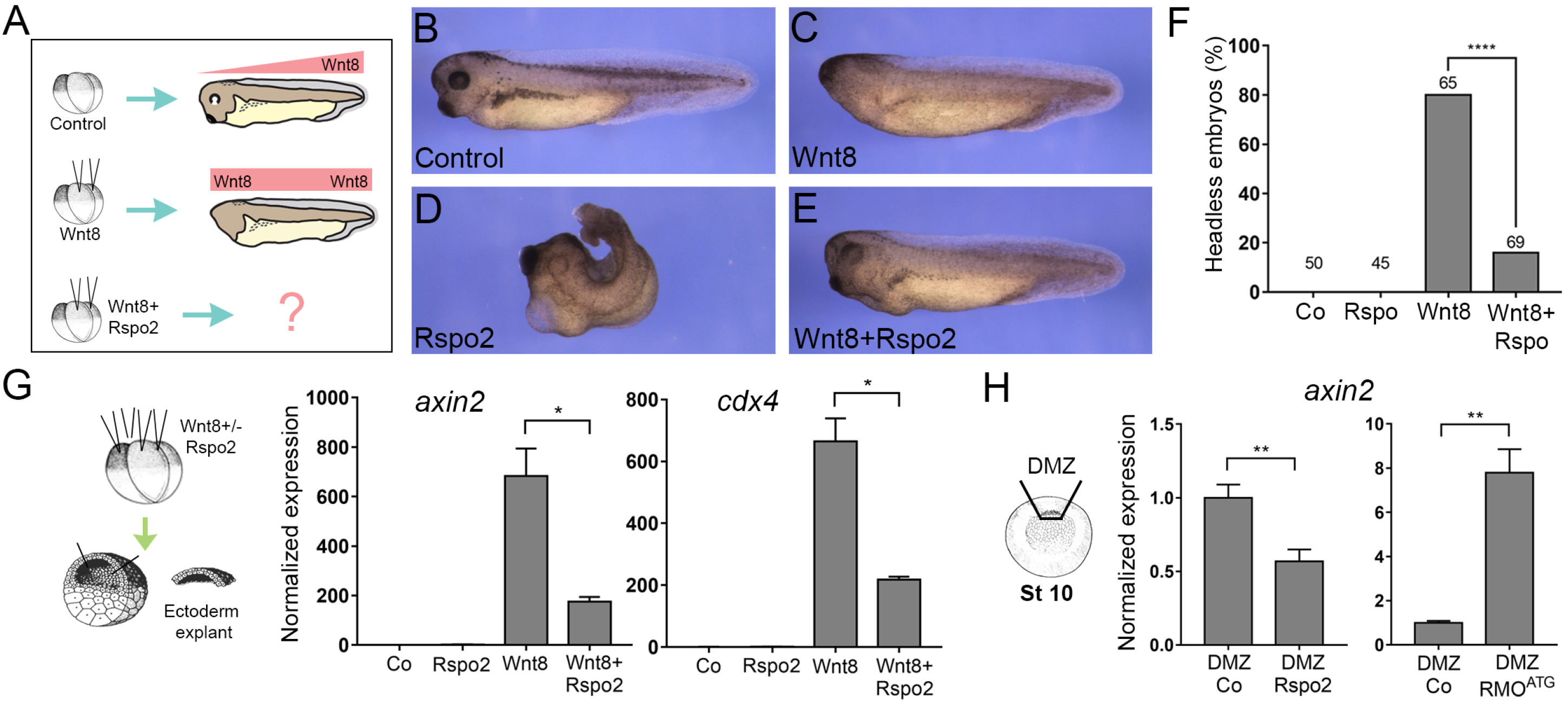
Rspo2 antagonizes Wnt signaling. (A) Scheme of the experiment. Four-cell embryos were injected animally into both dorsal blastomeres with the indicated constructs and cultured to stage 38. B, Uninjected control embryo. C, Headless embryo injected with Wnt8 DNA (50 pg). D, Embryo injected with Rspo2 RNA (0.5 ng). E, Embryo coexpressing Wnt8 DNA and Rspo2 mRNA. (F) Quantification of the data in A-D, representative of 3 independent experiments. Numbers of embryos per group are shown above each bar. ****, p<0.0001, Fisher’s exact test. (G) Target gene expression in Wnt8 and Rspo2-stimulated ectoderm explants. Embryos were injected into four animal blastomeres with Wnt8 DNA (50 pg) and Rspo2 RNA (0.5 ng), as indicated, and ectoderm explants were prepared at stage 9 and cultured until stage 13. (H) Dorsal marginal zones (DMZ) were dissected at stage 10 from the control and Rspo2 RNA- or RMO^ATG^-injected embryos and cultured until stage 12. (G, H) RT-qPCR analysis was carried out in triplicates for *axin2* and *cdx4*, and normalized to *eef1a1* levels. Means +/- s.d. are shown. Graphs are representative of three independent experiments. Statistical significance has been assessed by Student’s *t* test, *, p<0.05; **, p<0.01.

This result suggests that Rspo2 prevents the activation of specific Wnt target genes that are involved in posterior development (Ding et al., 2017; Kjolby and Harland, 2017; Nakamura et al., 2016; Nakamura and Hoppler, 2017). In ectoderm explants stimulated with Wnt8, Rspo2 downregulated the known Wnt targets *axin2* (Jho et al., 2002), *cdx4* (Northrop and Kimelman, 1994), *mesogenin1/msgn1* (Chalamalasetty et al., 2014; Wittler et al., 2007) and *myod1* (Hoppler et al., 1996) (Fig. 2G and Supplementary Fig. 3). Importantly, *axin2* was also inhibited by Rspo2 overexpression and upregulated after Rspo2 depletion in the marginal zone, where endogenous Wnt signaling takes place (Fig. 2H).

To confirm the specific effect of Rspo on Wnt signaling, we used the transgenic frog line *Xla. Tg(WntREs:dEGFP)^Vlemx^*, that contains a multimerized Wnt response element driving the expression of destabilized GFP (Tran et al., 2010). Coinjection of Rspo2 RNA into the transgenic embryos with mRFP RNA as a lineage tracer suppressed GFP fluorescence at the injected side (Fig. 3A-C), demonstrating the Wnt inhibitory activity of Rspo2.

**Figure 3.**
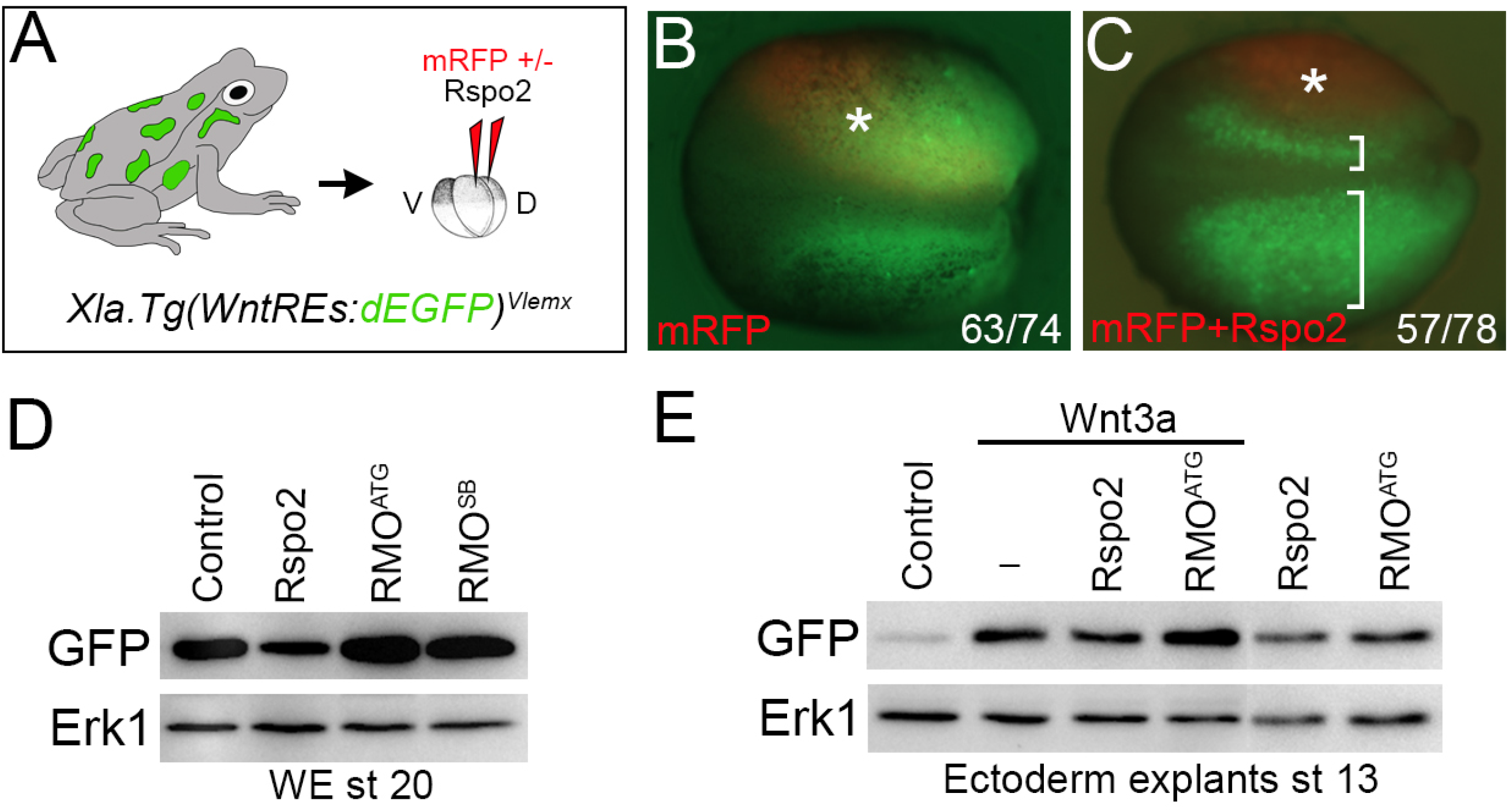
The effects of Rspo2 manipulation on Wnt reporter activity in transgenic embryos. (A) Experimental scheme. *Xla.Tg(WntREs:dEGFP)^Vlemx^* embryos were injected into one dorsal blastomere with mRFP RNA (50 pg) with (C) or without (B) Rspo2 RNA (0.5 ng). GFP fluorescence of the injected embryos at stage 18 is shown. Embryo images are representative of 3 different experiments. Asterisk indicates the injected side of the embryo, brackets in C show the comparison of the injected and the control sides. (D, E) Rspo2 modulates Wnt reporter activation. (D) Rspo2 RNA (0.5 ng), RMO^ATG^ (10 ng) or RMO^SB^ (20 ng) were injected at two dorsal blastomeres at 4-cell stage. The embryos were lysed at stage 20 and immunoblotted with anti-GFP antibodies. (E) Four-cell stage embryos were injected animally with Wnt3a RNA (50 pg) and Rspo2 RNAs (0.5 ng) or RMO^ATG^ (10 ng). Ectoderm explants were dissected at stage 9 and cultured until stage 13, then lysed and immunoblotted with anti-GFP antibodies. Erk1 is a control for loading in C, D.

This effect was estimated in a more quantitative way by immunoblotting of the lysates from both Rspo2-overexpressing and Rspo2-depleted embryos. Whole lysates from the embryos injected with Rspo2 RNA contained less GFP, whereas the lysates from the embryos injected with either MO contained more GFP, compared to the control embryos (Fig. 3D). Moreover, in ectoderm explants, Wnt3a-stimulated reporter activity was decreased by Rspo2 and upregulated by RMO^ATG^ (Fig. 3E).

Together, these findings indicate that Rspo2 antagonizes the Wnt pathway during anteroposterior axis specification.

### Rspo2 inhibits TCF3 phosphorylation

R-spondins are composed of two furin-like domains at the N-terminus, one thrombospondin type 1 domain (TSP) and the C-terminus enriched in basic amino acid residues (de Lau et al., 2014; Raslan and Yoon, 2019). To examine the mechanism, by which Rspo2 affects Wnt signaling, we assessed the ability of several Rspo2 constructs with specific domain deletions (Fig. 4A) to interfere with Wnt signaling.

**Figure 4.**
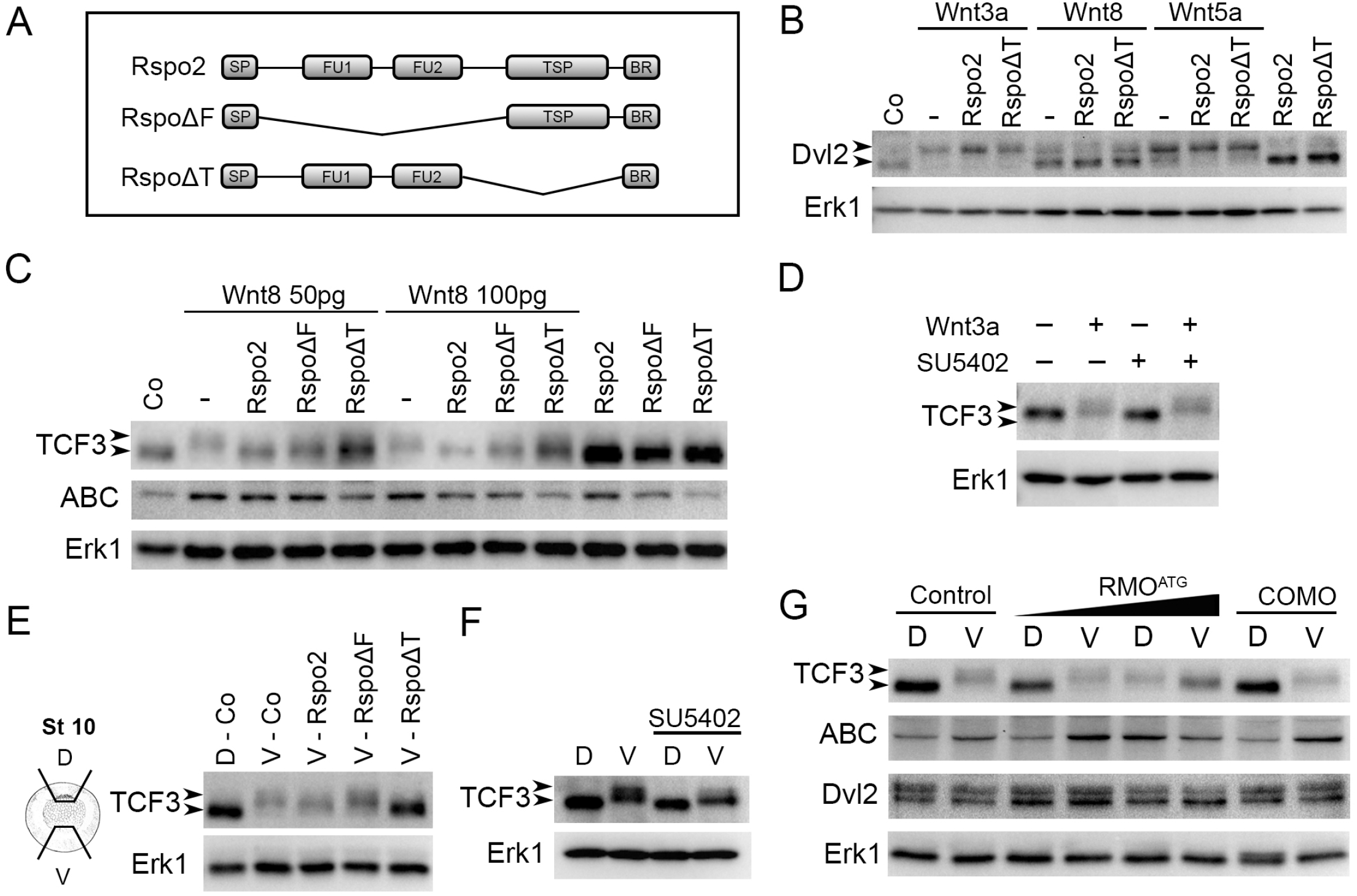
Rspo2 inhibits TCF3 phosphorylation. (A) Schematic of Rspo2 deletion constructs. SP, signal peptide; FU1, furin-like domain 1; FU2, furin-like domain 2; TSP, thrombospondin type 1 domain; BR, the basic amino acid-rich domain. (B, C) Effects of Rspo2 constructs on Wnt-dependent Dvl2 phosphorylation (B) and TCF3 phosphorylation and β-catenin levels (C). Four-cell stage embryos were injected animally with Wnt8 DNA (50 pg or 100 pg) or Wnt8, Wnt3a or Wnt5a RNAs (1 ng each) and Rspo2, RspoΔF or RspoΔT RNAs (0.5 ng each) as indicated. Ectoderm explants were dissected at stage 9 and cultured until stage 12 for immunoblotting with antibodies against Dvl2, TCF3, ABC (non-phosphorylated β-catenin). Arrowheads indicate the position of phosphorylated (upshifted) and non-phosphorylated Dvl2 or TCF3 proteins. Erk1 controls for loading. D, SU5402 does not block TCF3 phosphorylation in ectoderm stimulated by Wnt3a. E, Effects of Rspo2 constructs (0.5 ng each) on TCF3 phosphorylated by endogenous signals. Dorsal marginal zone (D) and ventral marginal zone (V) were dissected from the control and injected embryos at stage 10 and cultured until stage 12.5 for immunoblotting with anti-TCF3 antibodies as shown. F, Effects of SU5402 on TCF3 phosphorylation in marginal zone explants. G, Effects of Rspo2 depletion on TCF3 phosphorylation by endogenous signals. DMZ and VMZ explants of embryos injected with control MO (COMO, 20 ng) or RMO^ATG^ (20 ng) were dissected and analyzed by immunoblotting as in (B, C).

First, we asked which constructs retain the ability of full length Rspo2 to anteriorize the embryo. Rspo2 lacking the TSP domain (RspoΔT) had a strong cement gland-inducing activity (Supplementary Fig. 4A, D). RspoΔF also slightly enhanced head development, but the effect was much weaker than that of RspoΔT and the wild-type Rspo2 (Supplementary Fig. 4B, C). These observations are consistent with furin-like domains playing an important role in the inhibition of the Wnt pathway that is independent of the known Rspo2 receptors.

To address a specific mechanism of Wnt pathway inhibition by Rspo2, we analyzed Dvl phosphorylation, a common proximal event in Wnt/Frizzled signaling (Angers and Moon, 2009; Yanagawa et al., 1995). Phosphorylated Dvl2 migrated slower in ectoderm cells stimulated with Wnt8, Wnt3a and Wnt5a. Rspo2 constructs did not affect Dvl2 mobility on their own or in response to Wnt signals (Fig. 4B), suggesting that Rspo2 does not operate by modulating the activity of Wnt ligands or Frizzled receptors.

We next evaluated the effect of Rspo2 on the downstream signaling intermediates TCF3 and β-catenin. The phosphorylation of TCF3 in response to a Wnt signal leads to TCF3 dissociation from target promoters and transcriptional derepression of Wnt target genes (Hikasa et al., 2010). TCF3 phosphorylation was visualized by the slower mobility of the TCF3 band from the lysates of ectoderm explants expressing Wnt8 (Fig. 4C). TCF3 migrated faster in the lysates of cells co-expressing Rspo2. Importantly, both RspoΔF and RspoΔT inhibited TCF3 phosphorylation, although RspoΔF was less effective in this assay. In the absence of Wnt ligands, we observed that TCF3 levels were consistently higher in cells expressing Rspo2 constructs, suggesting that Rspo2 might also influence TCF3 protein stability. In the same experiment, levels of non-phosphorylated β-catenin increased in response to Wnt8, and this effect was reversed by Rspo2 constructs (Fig. 4C).

Many Wnt gene targets are also controlled by the FGF pathway (Kjolby et al., 2019; McGrew et al., 1997) and the FGF pathway was reported essential for Wnt activity during anteroposterior patterning (Domingos et al., 2001). Since Rspo2 reduces FGF signaling in the same system (Reis and Sokol, 2020), it is possible that the effect of Rspo constructs on TCF3 is indirect, due to the suppression of the FGF pathway. Treating the explants with the FGF receptor inhibitor SU5402 under the conditions when FGF signaling is completely blocked (Fletcher and Harland, 2008; Mohammadi et al., 1997) did not interfere with TCF3 phosphorylation in response to Wnt3a (Fig. 4D). This result indicates that TCF3 phosphorylation by Wnt3a does not require FGF signaling and that the effect of Rspo2 constructs on the Wnt pathway is direct, rather than indirect, through the suppression of FGF signaling activity.

Our conclusions have been extended to endogenous Wnt signaling that is responsible for TCF3 phosphorylation in the mesoderm (marginal zone) during gastrulation (Hikasa et al., 2010). Rspo2 and RspoΔT constructs inhibited TCF3 phosphorylation in ventral marginal zone explants, while RspoΔF had a mild effect (Fig. 4E). Notably, SU5402 had little effect on TCF3 phosphorylation in these explants, further indicating that TCF3 is regulated predominantly by the Wnt pathway (Fig. 4F). These observations support our conclusion that Rspo2 antagonizes Wnt signaling by blocking TCF3 phosphorylation.

Furthermore, TCF3 phosphorylation became prominent in the dorsal marginal zone explants isolated from Rspo2 morphants (Fig. 4G). This effect correlated with the accumulation of non-phosphorylated β-catenin. By contrast, no significant changes in Dvl2 levels or mobility have been observed, suggesting that Frizzled receptors are not involved. Based on these results, we propose that Rspo2 enhances anterior development by inhibiting TCF3 phosphorylation.

### The Wnt-inhibitory activity of Rspo2 relies on TCF3

If Rspo2 modulates Wnt target genes by inhibiting TCF3 phosphorylation, the depletion of TCF3 should prevent Rspo2 gain-of-function phenotype. Consistent with this prediction, the anteriorized phenotype of RspoΔT-expressing embryos was suppressed by TCF3 depletion (Fig. 5A, B). RspoΔT protein levels did not change in TCF3-depleted embryos, supporting knockdown specificity (Fig. 5C). This result suggests that the Wnt antagonistic activity of Rspo2 requires TCF3.

**Figure 5.**
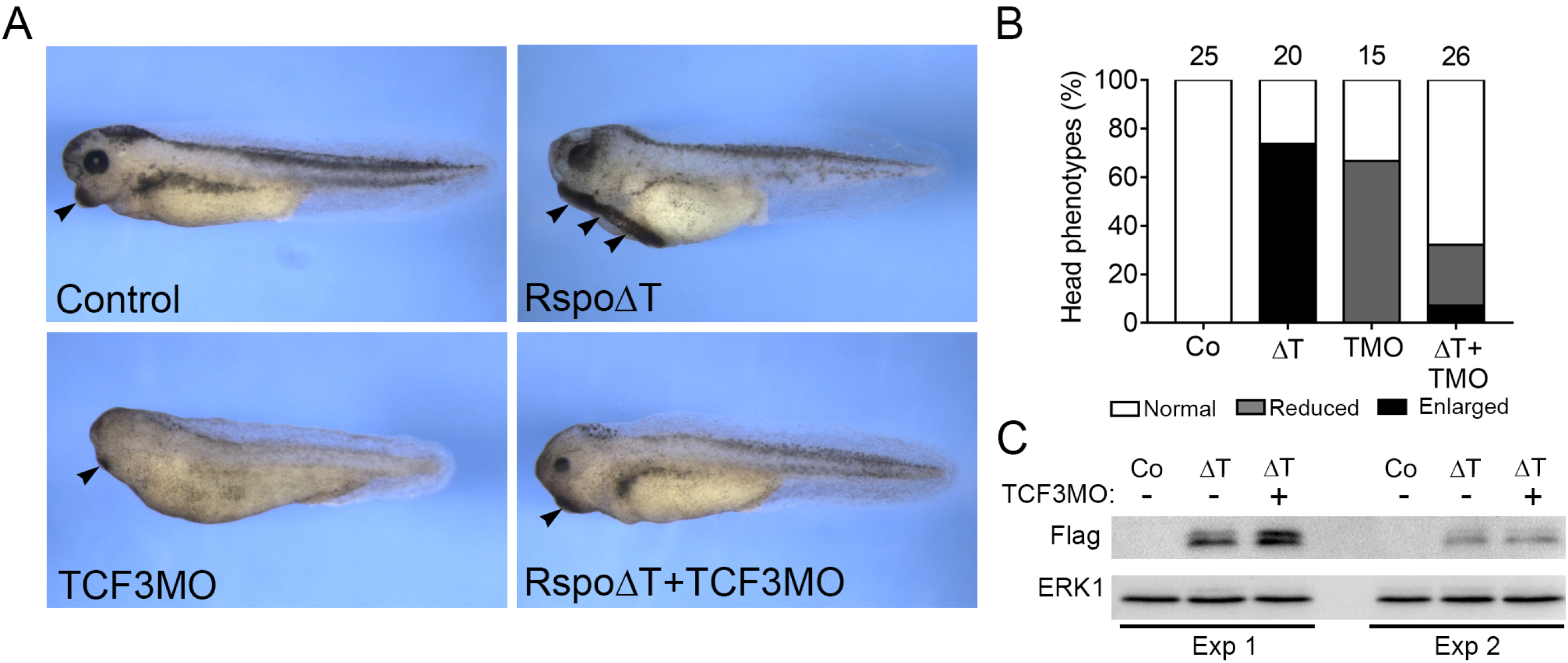
TCF3 is essential for Rspo2 inhibitory effects. A, TCF3MO rescues the anteriorized phenotype of RspoΔT RNA overexpressing embryos. Four-cell stage embryos were dorsally injected with TCF3MO (30 ng) and/or RspoΔT RNA (0.5 ng). Arrowheads indicate the cement gland. B, Quantification of the data in A, representative of two independent experiments. Numbers of embryos per group are shown above each bar. C, RspoΔT expression levels are not altered by TCF3MO in ectoderm explants (stage 12) in two independent experiments (Exp 1 and Exp 2). ΔT, RspoΔT; TMO, TCF3MO.

In a converse experiment, Rspo2 depletion is predicted to be rescued by a constitutive TCF3 repressor that does not bind β-catenin (ΔβTCF3) (Hikasa et al., 2010). Supporting this expectation, the effect of Rspo2 depletion on both anterior (*otx2* and *ag1*), and posterior (*cdx4* and *msgn1*) markers were partially rescued in the morphants by ΔβTCF3 (Fig. 6A).

**Figure 6.**
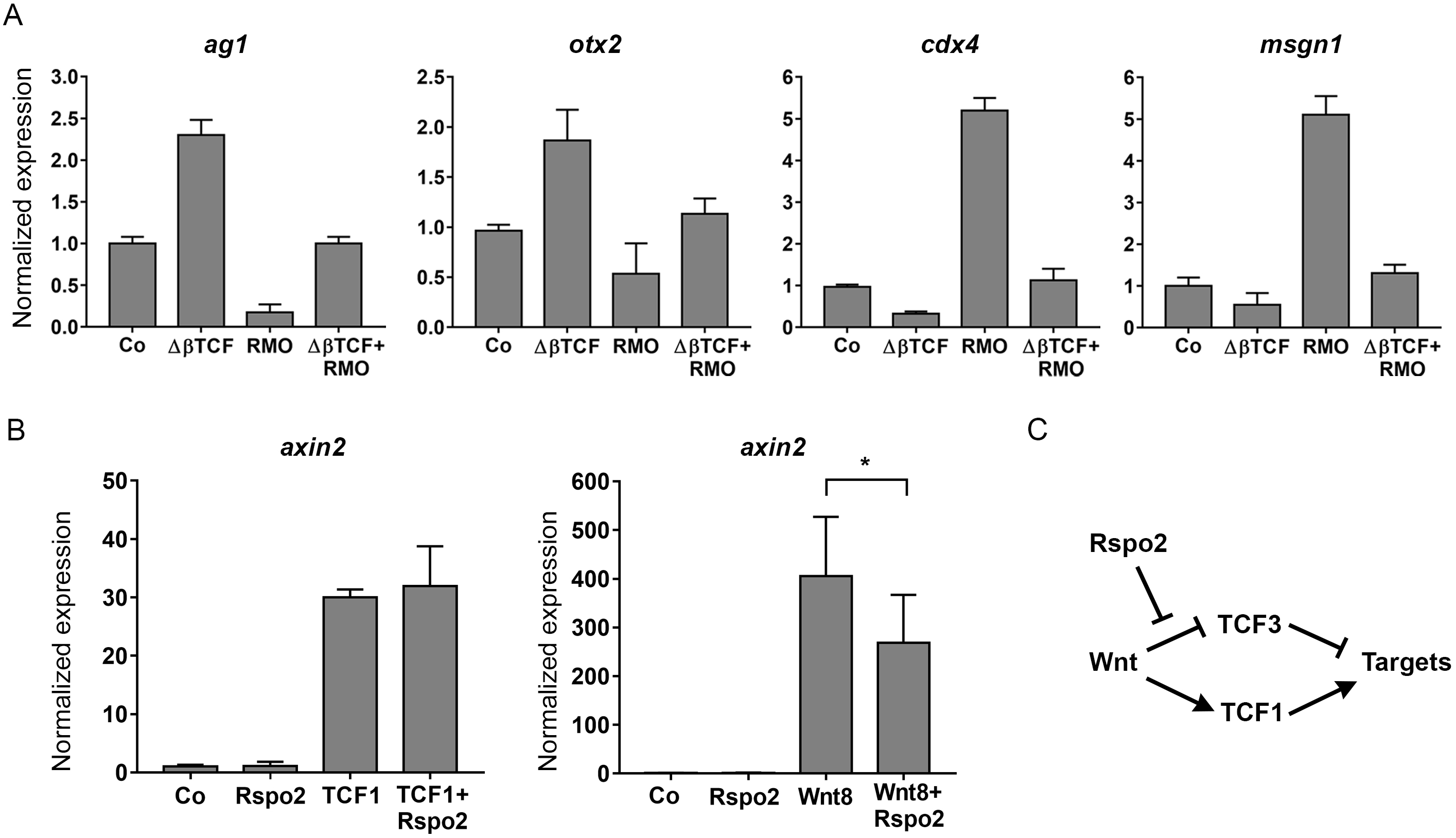
Rspo2 inhibits Wnt signaling through TCF3. A, ΔβTCF3 RNA (10 pg) rescues *ag1, otx2, cdx4*, and *msgn1* expression in embryos injected with 10 ng of RMO^ATG^. B, Rspo2 inhibits *axin2* upregulation by Wnt8 but not TCF1 in ectoderm cells. Embryos were injected with Wnt8 (20 pg) or TCF1 (100 pg) RNA without or with Rspo2 RNA (300 pg). Ectoderm explants were prepared at stage 8.5-9 and analyzed at stage 13. RT-qPCR analysis was carried out in triplicates for *axin2* and normalized to *eef1a1* levels. Means +/- s.d. are shown. Graphs are representative of 2-4 independent experiments. Statistical significance has been assessed by Student’s *t* test, *, p<0.05. C, Model for Rspo2-mediated repression. Rspo2 inhibits Wnt target activation mediated by TCF3 phosphorylation but not the TCF1-dependent response.

Based on these observations, we propose that the Wnt-inhibitory function of Rspo2 is mediated by TCF3, a predominant TCF in early embryos that functions as a transcriptional repressor. By contrast, other TCF proteins mediating Wnt signaling, such as TCF1/Tcf7 or Lef1, can activate Wnt targets. Notably, Rspo2 did not downregulate *axin2* induction by *tcf1* RNA in ectoderm cells, whereas it significantly reduced Wnt8 activity in the same experiment (Fig. 6B). This observation suggests a model, in which Rspo2 prevents the ability of Wnt signaling to inhibit TCF3 repressive activity, but does not downregulate TCF1-dependent signaling (Fig. 6C).

## Discussion

This study has been focused on Rspo2, a member of the R-spondin family of Wnt pathway modulators. We demonstrate that Rspo2 promotes anterior development by inhinbiting TCF3 phosphorylation and Wnt target genes activation independently of the known interaction with LGR4/5 and ZNRF3/RNF43 receptors. In addition to the Wnt pathway, R-spondins were described to affect TGFβ (Kazanskaya et al., 2004; Zhou et al., 2017) and FGF (Reis and Sokol, 2020; Zhang et al., 2017) signaling. Although R-spondins are well known to potentiate Wnt signals in various cells and embryonic tissues (de Lau et al., 2014; Kazanskaya et al., 2004; Kim et al., 2008; Nam et al., 2006; Raslan and Yoon, 2019; Wei et al., 2007), we demonstrate an alternative role for Rspo2 as a Wnt antagonist during anteroposterior patterning. While surprising, this conclusion is consistent with other reports using zebrafish and cancer cell lines (Rong et al., 2014; Wang et al., 2018; Wu et al., 2014). We propose that Rspo2 modulates the Wnt pathway in a context-specific manner.

Similar to other secreted multidomain molecules, Rspo2 is a pleiotropic regulator of signaling. We find that Rspo2 inhibits the Wnt pathway via the Furin-like and the TSP domains, however, it antagonizes the FGF pathway exclusively via the TSP domain (Reis and Sokol, 2020). The binding of LGR4/5 and ZNRF3/RNF43 receptors to the furin-like domains does not seem to be involved in the Wnt inhibitory activity of Rspo2. The effect of the TSP domain could be mediated by its interactions with heparan sulfate proteoglycans that are known to modulate both FGF and Wnt signaling (Lin and Perrimon, 1999; Rapraeger et al., 1991; Yayon et al., 1991). These experiments further illustrate the complexity of the Wnt-FGF crosstalk extending from the extracellular level (Yamamoto et al., 2005) to transcriptional regulation (Haremaki et al., 2003; Kjolby et al., 2019).

The main mechanism for R-spondin signaling in adult stem cells is the modulation of Frizzled degradation by the interaction with LGR4/5 and ZNRF3/RNF43 receptors (de Lau et al., 2011; Glinka et al., 2011; Hao et al., 2012). The phosphorylation of Dishevelled, a proximal marker of Wnt/Frizzled signaling, was not altered in embryonic tissues with manipulated Rspo2 function. This finding suggests that Frizzled signaling is not involved. Moreover, mutations abolishing the binding of LGR4/5 and ZNRF3/RNF43 did not affect the anteriorizing activity of Rspo2, indicating that these interactions are not involved. At present, we cannot exclude a role for LRP5/6 in mediating Rspo2 function, as it was reported to interact with Rspo1, a closely related protein (Binnerts et al., 2007; Wei et al., 2007). Consistent with recent reports (Dubey et al., 2020; Lebensohn and Rohatgi, 2018; Park et al., 2018; Szenker-Ravi et al., 2018), we propose that Rspo2 is a context-dependent Wnt antagonist that may function via yet unknown receptors.

Whereas the Rspo2 receptors mediating its effects on early embryos are not known, we present mechanistic evidence that Rspo2 functions by inhibiting TCF3 phosphorylation. TCF3 is a transcriptional repressor of Wnt targets that is inactivated by Wnt-dependent phosphorylation during anteroposterior patterning (Hikasa et al., 2010). This phosphorylation is blocked by Rspo2, thereby preventing Wnt target activation. In support of this conclusion, non-phosphorylatable TCF3 rescued Wnt target gene expression in Rspo2-depleted embryos. It is currently unknown whether Rspo2 modulates the phosphorylation of other TCF proteins, including the ones with a positive effect on transcription, such as TCF1 (Cadigan and Waterman, 2012; Sokol, 2011). Notably, Rspo2 did not inhibit the activity of TCF1 in our experiments. In a different developmental context, in which the TCF3 is not expressed, yet another TCF protein is phosphorylated by a Wnt signal (Adam et al., 2018; Hikasa and Sokol, 2011), R-spondins might potentiate Wnt signaling through the same mechanism. Several TCF proteins are known to be phosphorylated by HIPK2, Nemo-like kinase and casein kinases 1 and 2 (Hammerlein et al., 2005; Hikasa and Sokol, 2011; Ota et al., 2012; Smit et al., 2004), but upstream pathways leading to the activation of these protein kinases remain to be clarified. Additional work is needed to fully understand the molecular basis for the context-dependent activity of Rspo2 in embryonic development.

## Methods

### Plasmids, in vitro RNA synthesis and morpholino oligonucleotides (MOs)

The DNA clone 6988843 encoding *X. tropicalis* Rspo2 was obtained from Dharmacon. The plasmid encoding full length Rspo2 (pCS2-Rspo2-Flag) was generated by inserting PCR-amplified coding region of Rspo2 into the EcoRI and BamHI sites of pCS2-Flag. Various Rspo2 constructs (Supplementary Table 2) were generated using single primer-based site-directed mutagenesis as described (Itoh et al., 1995). pCS2-RspoΔF-Flag lacks amino acids 37-134. pCS2-RspoΔT-Flag lacks amino acids 147-204. Alanine substitutions have been made in pCS2-Rspo2-Flag or pCS2-RspoΔT-Flag in the furin-like domain 1 (R65A or Q70A), and furin-like domain 2 (F105A or F109A) to generate Rspo2 that does not bind ZNRF3/RNF43 or LGR4/5 as described (Xie et al., 2013). All constructs were verified by Sanger sequencing. Details of cloning are available upon request.

Capped mRNAs were synthesized using mMessage mMachine kit (Ambion, Austin, TX). The following linearized plasmids have been used as templates: pSP64T-Wnt3a (Wolda et al., 1993), pSP64T-Wnt8 (Christian et al., 1991), pCS2-Wnt8, and pSP64T-Wnt5a (Moon et al., 1993), ΔβTCF3 (Hikasa et al, 2010), pCS2-TCF1 (Hikasa and Sokol, 2011), pCS2-mRFP (membrane-targeted), pCS2-Rspo-Flag, pCS2-RspoΔF, pCS2-RspoΔT, pCS2-RspoR65A-Flag, pCS2-RspoQ70A-Flag, pCS2-RspoF105A-Flag, and pCS2-RspoF109A-Flag. The following MOs have been purchased from Gene Tools (Philomath, OR): RMO^ATG^, 5’-AAAGAGTTGAAACTGCATTTGG-3’, RMO^SB^, 5’-GCAGCCTGGATACACAGAAACAAGA-3’, control MO (CoMO), 5’-GCTTCAGCTAGTGACACATGCAT-3’. TCF3MO has been described previously (Hikasa et al., 2010).

### Xenopus embryo culture, microinjections, imaging and statistical analysis

*In vitro* fertilization and culture of *Xenopus laevis* embryos were carried out as described (Dollar et al., 2005). Staging was according to Nieuwkoop and Faber (Nieuwkoop and Faber, 1967). Wnt reporter pbin7LefdGFP transgenic embryos (Tran et al., 2010) have been obtained from the National Xenopus Resource (Woods Hole, MA). For microinjections, four-cell embryos were transferred into 3 % Ficoll in 0.5x Marc’s Modified Ringer’s (MMR) buffer (50 mM NaCl, 1 mM KCl, 1 mM CaCl_2_, 0.5 mM MgCl_2_, 2.5 mM HEPES pH 7.4) (Peng, 1991) and 10 nl of mRNA or MO solution was injected into one or more blastomeres. Amounts of injected mRNA and MOs per embryo, indicated in figure legends, have been optimized in preliminary dose-response experiments. Control MO was injected as at a dose that matched the highest amount of any other MO used in the same experiment.

Embryos were imaged at the indicated stages using Leica Wild M10 stereomicroscope using the OpenLab software. Unless otherwise specified, each experiment has been carried out at least three times. Statistical analyses were performed using GraphPad Prism 6 software. Data are mean±s.d. and statistical significance was assessed using an unpaired two-tailed Student’s t-test or Fisher’s exact test. Significant differences are indicated by p values, e. g. *, p<0.05; **, p<0.01; ****, p<0.0001.

### Ectoderm and marginal zone explants, RNA sequencing, RT-qPCR

Ectoderm explants were prepared at late blastula stages and cultured until the indicated time to observe morphological changes or lysed for RNA extraction or immunoblotting. Marginal zone explants were dissected at early gastrula stage and cultured until stage 12.5 when they were lysed for immunoblot analysis. To inhibit FGF receptor activity, ectoderm explants or marginal zone explants have been cultured with SU5402 (100 μM, Calbiochem) from the time of isolation until they were lysed for immunoblot analysis.

For quantitative PCR (RT-qPCR) and RNA sequencing, RNA was extracted from a group of 4-5 embryos, ten animal caps or ten marginal zone explants, at stages 10 or 12.5, using RNeasy kit (Qiagen). RNA sequencing was carried out using the HiSeq PE150 platform (150 b.p., paired end sequencing) and analyzed by Novogene (Sacramento, CA). cDNA was made from 1 μg of total RNA using iScript (Bio-Rad). qPCR reactions were amplified using a CFX96 light cycler (Bio-Rad) with Universal SYBR Green Supermix (Bio-Rad). Primer sequences used for RT-qPCR are listed in Supplementary Table 2. Data represent at least 3 independent experiments each including triplicate samples. All samples were normalized to control embryos. *eef1a1* served as an internal control. Means +/- s. d. are shown. Statistical significance was assessed using the Student’s *t*-test.

### Immunoblot analysis

Immunoblot analysis was carried out essentially as described (Itoh et al., 2005). Briefly, 10 animal caps or 7 marginal zone explants at stage 12.5 were homogenized in 50 μl of the lysis buffer (50 mM Tris-HCl pH 7.6, 50 mM NaCl, 1 mM EDTA, 1% Triton X-100, 10 mM NaF, 1 mM Na_3_VO_4_, 25 mM β-glycerol phosphate, 1 mM PMSF). After centrifugation for 3 min at 16000 g, the supernatant was subjected to SDS-PAGE and western blot analysis following standard protocols (Itoh et al., 2005). The following primary antibodies were used: mouse anti-FLAG (M2, Sigma), mouse anti-non-phosphorylated β-catenin (ABC; Upstate Biotechnology), rabbit anti-XTCF3N (Zhang et al., 2003), rabbit anti-Dvl2 (Itoh et al., 2005). Staining with rabbit anti-Erk1 (Cell Signaling) was used as loading control. Chemiluminescence was captured by the ChemiDoc MP imager (BioRad).

### In situ hybridization

Whole-mount in situ hybridization with the digoxigenin-labeled antisense RNA probes for *krt12.4* (Winkles et al., 1985), *foxg1/BF1* (Bourguignon et al., 1998), and *cdx4* (Reis and Sokol, 2020), was carried out as described (Harland, 1991).

## Acknowledgements

We thank Miho, Matsuda, Keiji Itoh and Jean-Pierre Saint-Jeannet for the comments on the manuscript. We also thank Aurelian Radu for the help with the analysis of RNA sequencing, Olga Ossipova for qPCR primers, Pamela Mancini for advice on Adobe Illustrator and members of the Sokol laboratory for discussions. This study has been supported by the NIH grant HD092990 to SYS.

## Author contributions

A.H.R. designed experiments, carried out experiments, analyzed data and wrote the manuscript. S.Y.S. designed experiments, analyzed data and wrote the manuscript.

## Notes

### Competing Interest Statement

The authors have declared no competing interest.

