## Supplementary material for "Rspo2 inhibits TCF3 phosphorylation to antagonize Wnt signaling during vertebrate anteroposterior axis specification": suppl data

**Supplementary Information:**

**Supplementary Figure legends**

**Supplementary Fig. 1: Top target genes upregulated by Rspo2 in ectoderm**. Top ten target genes that encode transcription factors are shown. Four-cell embryos were injected with 0.5 ng of Rspo RNA into animal blastomeres. Ectoderm explants were prepared from injected embryos at stage 9, RNA was isolated at stage 10 for RNA-sequencing. Human gene symbols are shown, except for those that are *Xenopus*-specific (*). Target genes of the injected animal caps have been normalized to the uninjected controls.

**Supplementary Fig. 2: The binding of ZNRF3/RNF43 and LGR4/5 is not required for the ability of Rspo2 to anteriorize the embryo**. A, Scheme of the point mutant constructs generated from the full length Rspo2. B-E, Cement gland enlargement (dashed white line) caused by Rspo2 RNA injection. Ventral animal blastomeres of four-cell embryos were injected with 500 pg of Rspo2, R65A, Q70A, F105A or F109A mRNA. F, Quantification of the embryos showing enlarged cement gland after overexpression of different Rspo2 constructs. Numbers of embryos per group are shown on the top of each bar.

**Supplementary Fig. 3: Rspo2 inhibits Wnt target genes in ectoderm explants.** Wnt target gene expression in ectoderm explants at stage 13. Embryos were injected into four animal blastomeres with Wnt8 DNA (50 pg) and Rspo2 RNA (0.5 ng). RT-qPCR analysis was carried out in triplicates for *myod1* and *msgn1*.

**Supplementary Fig. 4. Comparison of the anteriorizing activity of Rspo2 constructs.** Ventral animal blastomeres of four-cell embryos were injected with 500 pg of RNA encoding Rspo2 deletion mutants. Embryo images, representative of three independent experiments, are shown. The ratio of the number of anteriorized embryos to the total number of injected embryos is indicated. A, Control embryo at stage 27. B-D, the sibling embryos injected with RNAs encoding full-length Rspo2 (B), Rspo∆F (C), Rspo∆T (D).

**Supplementary Tables 1-2.**

**Supplementary Table 1.** Quantification of data for wholemount in situ hybridization shown in representative images of Figure 1E-J. Numbers of embryos with indicated changes in gene expression are indicated.

**Supplementary Table 2.** Primer sequences for the site-directed mutagenesis and RT-qPCR.

| Top genes upregulated by Rspo2 | | |
|---|---|---|
| Human symbol | log2(Fold_change) normalized | p-value |
| OTX2 | 5,20 | 1,0882E-147 |
| TCF24 | 4,70 | 0,00023719 |
| CRX/otx5* | 3,98 | 2,7623E-183 |
| GBX2 | 3,80 | 0,010773101 |
| ZIC3 | 3,61 | 4,00229E-35 |
| NKX2-8 | 3,46 | 0,002037888 |
| KLF5 | 3,46 | 0,002037888 |
| hes3.L* | 3,32 | 0,040847184 |
| hes5.1.L* | 3,32 | 0,003825822 |
| NFKB1 | 3,32 | 0,040847184 |

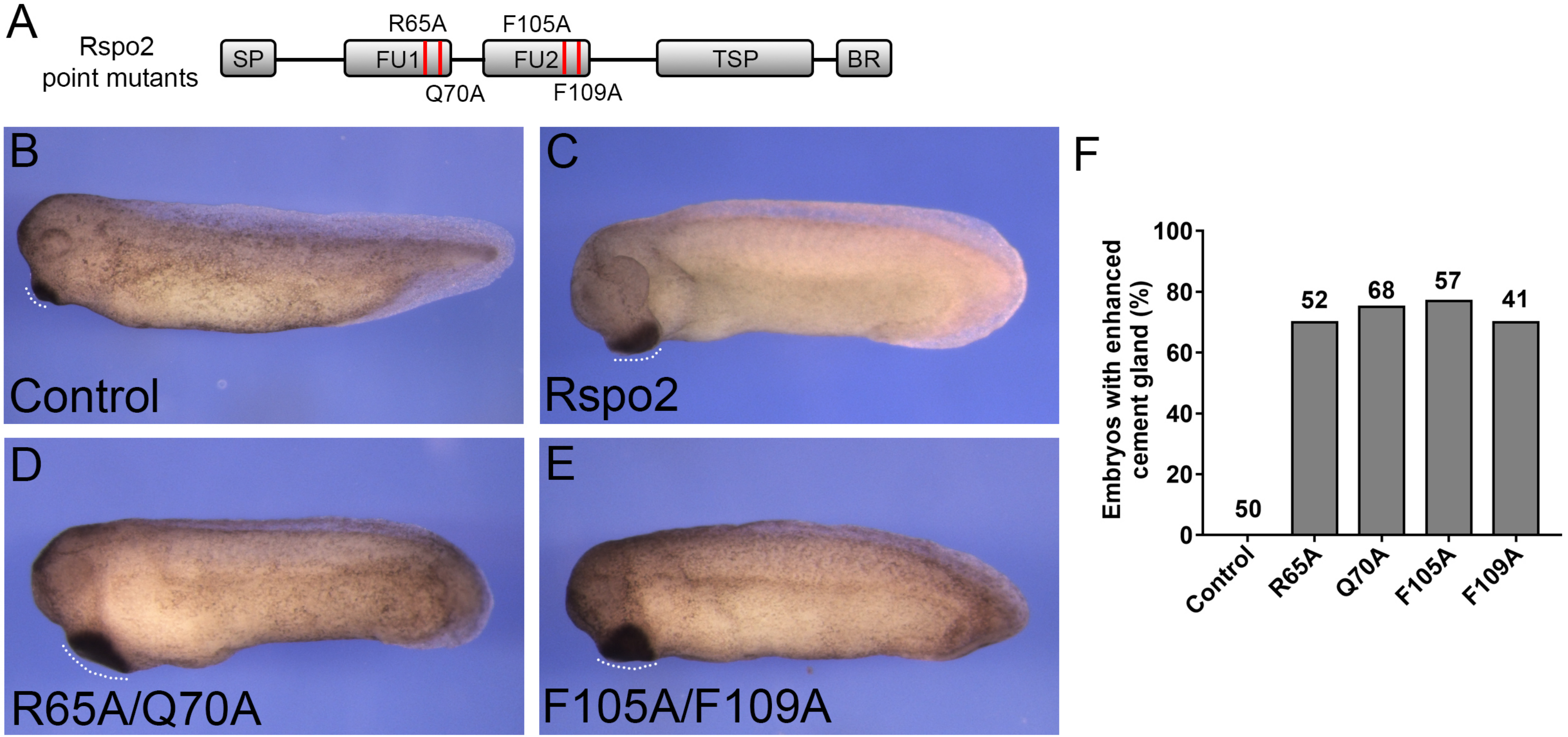
A
Rspo2
point mutants
R65A
F105A
SP
FU1
FU2
TSP
BR
Q70A
F109A
B
Control
C
Rspo2
D
R65A/Q70A
E
F105A/F109A
F
Embryos with enhanced cement gland (%)
100
80
60
40
20
0
50
52
68
57
41
Control
R65A
Q70A
F105A
F109A

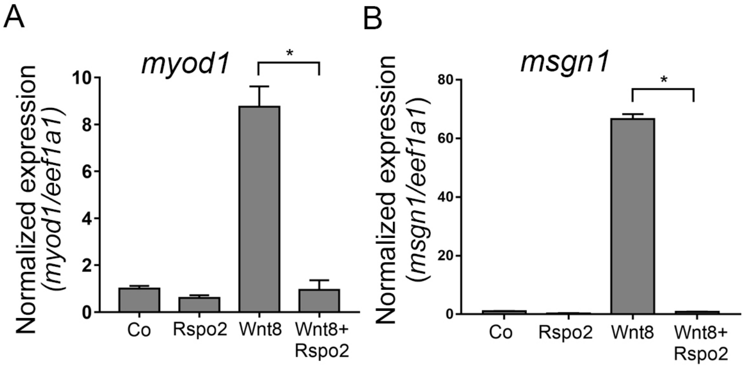
A
myod1
*
Normalized expression
(myod1/eef1a1)
10
8
6
4
2
0
Co
Rspo2
Wnt8
Wnt8+
Rspo2
B
msgn1
*
Normalized expression
(msgn1/eef1a1)
80
60
40
20
0
Co
Rspo2
Wnt8
Wnt8+
Rspo2

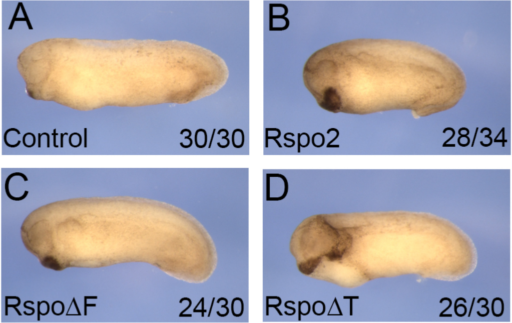
A
Control
30/30
B
Rspo2
28/34
C
RspoΔF
24/30
D
RspoΔT
26/30

**Quantification of in situ hybridization analysis in embryos with modulated Rspo2 levels**

| ***krt12.4*** | **Co** | **Rspo** | **ATGMO** |
|---|---|---|---|
| Increased | 0 | 0 | 13 |
| Decreased | 0 | 7 | 0 |
| Unchanged | 30 | 3 | 4 |

| ***foxg1*** | **Co** | **Rspo** | **ATGMO** |
|---|---|---|---|
| Increased | 0 | 5 | 0 |
| Decreased | 0 | 0 | 12 |
| Unchanged | 22 | 7 | 3 |

| ***cdx4*** | **Co** | **Rspo** | **ATGMO** |
|---|---|---|---|
| Increased | 3 | 0 | 12 |
| Decreased | 0 | 7 | 0 |
| Unchanged | 19 | 5 | 3 |

**Supplementary Table S2.**

**Primers for Rspo2 mutagenesis**

Rspo∆F:

5'- AGACGGAGCAAGAGAGCCAGATCTCCATTGGATGACACCATG-3'

Rspo∆T:

5' -TGCGTGGATGGCTGTGAAGCTAGCGGAGGAACAAGAACCACA-3'

R65A:

5'- ACTGTTTTTCTATCTGCGCGCAGAAGGTATGAGGCAGTAT-3'

Q70A:

5'-CGAAGAGAAGGTATGAGGGCATATGGAGAGTGTCTGCAG-3'

F105A:

5'-GAAAATTGTGACTCTTGTGCATGCCGAGATTTTTGCATAA-3'

F109A:

5'-TCTTGTTTTAGCCGAGATGCATGCATAAAGTGCAAATCT-3'

**Primers for RT-qPCR**

| | |
|---|---|
| *otx2.L*: | F: 5'-GGATGGATTTGTTACATCCGTC-3' |
| | R: 5'-CACTCTCCGAGCTCACTTCCC-3' |
| *ag1.S*: | F: 5'-GGTGCTGCCAAGTCTGAGC-3' |
| | R: 5'-GCCAGTTTCTGTGCCATTTTGTCA-3' |
| *krt12.4.L*: | F: 5'-CACCAGAACACAGAGTAC-3' |
| | R: 5'-CAACCTTCCCATCAACCA-3' |
| *cdx4.L*: | F: 5'-TGATTTATCACCTAACCAG-3' |
| | R: 5'-GTCCCAGATGGATGAGGAGA |
| *msgn1.L*: | F: 5'-GTATCCAACACTTTGCCATG-3' |
| | R: 5'-AGCACTGGAGAAGGTTTGTG-3' |
| *axin-2-like*: | F: 5'-GGCTGGTCTCTCTGCCTCTT-3' |
| | R: 5'-TGTCCTTCTCCTCCTGCTTCT-3' |
| *eef1a1.S*: | F: 5'-ACCCTCCTCTTGGTCGTTTT-3' |
| | R: 5'-TTTGGTTTTCGCTGCTTTCT-3' |
| *myod1*: | F: 5'-AGGTCCAACTGCTCCGACGGCATGAA-3' |
| | R: 5'-AGGAGAGAATCCAGTTGATGGAAACA-3' |
